# FSHR-1/GPCR activates the mitochondrial unfolded protein response in *Caenorhabditis elegans*

**DOI:** 10.1101/586883

**Authors:** Sungjin Kim, Derek Sieburth

## Abstract

The mitochondrial unfolded protein response (UPR^mt^) is an evolutionarily conserved adaptive response that functions to maintain mitochondrial homeostasis following mitochondrial damage. In *C. elegans*, the nervous system plays a central role in responding to mitochondrial stress by releasing endocrine signals that act upon distal tissues to activate the UPR^mt^. The mechanisms by which mitochondrial stress is sensed by neurons and transmitted to distal tissues is not fully understood. Here, we identify a role for the conserved follicle-stimulating hormone G protein coupled receptor (GPCR), FSHR-1, in promoting UPR^mt^ activation. Genetic deficiency of *fshr-1* severely attenuates UPR^mt^ activation and organism-wide survival in response to mitochondrial stress. FSHR-1 functions in a common genetic pathway with SPHK-1/sphingosine kinase to promote UPR^mt^ activation, and FSHR-1 regulates the mitochondrial association of SPHK-1 in the intestine. Through tissue-specific rescue assays, we show that FSHR-1 functions in neurons to activate the UPR^mt^, to promote mitochondrial association of SPHK-1 in the intestine, and to promote organism-wide survival in response to mitochondrial stress. We propose that FSHR-1 functions cell non-autonomously in neurons to activate UPR^mt^ upstream of SPHK-1 signaling in the intestine.

## Introduction

The mitochondrial unfolded protein response (UPR^mt^) functions to maintain mitochondrial protein homeostasis in response to mitochondrial dysfunction caused by mitochondrial DNA damage, incorrect mitochondrial protein folding, or impaired oxidative phosphorylation. Failure to appropriately control and maintain protein homeostasis in the mitochondria is associated with the development of numerous diseases, neurodegeneration, and ageing (Durieux *et al*. 2011; Liu *et al*. 2014; Pellegrino *et al*. 2014; Nargund *et al*. 2015; Fiorese *et al*. 2016; Martinez *et al*. 2017). The UPR^mt^ is initiated when mitochondrial proteostasis is disrupted, detection of which by mitochondrial and cytosolic factors leads to epigenetic modifications and transcriptional responses in the nucleus that restore mitochondrial function. A critical sensor and activator of the UPR^mt^ in *C. elegans* and in mammals is the leucine zipper transcription factor ATSF-1/ATF5 (Fiorese *et al*. 2016). ATFS-1 is normally targeted to mitochondria where it is degraded, but upon mitochondrial stress, mitochondrial import is disrupted and ATFS-1 is targeted instead to the nucleus where it regulates the expression of a cascade of genes including the conserved HSP70-like chaperone, *hsp-6* which is targeted to the mitochondria to restore protein folding (Nargund *et al*. 2012). The intestinal UPR^mt^ can be activated by mitochondrial stress originating either cell autonomously in the intestine or cell non-autonomously in the nervous system. Mitochondrial stress in neurons activates the UPR^mt^ in intestine through the release of neuropeptides, serotonin and/or Wnt ligands (Berendzen *et al*. 2016; Shao *et al*. 2016; Zhang *et al*. 2018).

Genetic screens for additional factors that activate the UPR^mt^ have revealed important roles for mitochondrial ceramide produced by SPTL-1/serine palmitoyltransferase and sphingolsine-1-phosphate (S1P) produced by SPHK-1/sphingosine kinase in the activation of the UPR^mt^ (Liu *et al*. 2014; Kim and Sieburth 2018a). SPHK-1 recruitment to mitochondria from cytoplasmic pools may serve as an early signal to activate UPR^mt^ (Kim and Sieburth 2018a). SPHK-1 mitochondrial recruitment is positively regulated by mitochondrial stress originating either from the intestine or from the nervous system. Neuronal mitochondrial stress activates intestinal SPHK-1 by a mechanism that involves neuropeptide but not serotonin signaling (Kim and Sieburth 2018a). Neuropeptides exert their biological functions primarily through activating G protein coupled receptors (GPCRs) on target cells to trigger downstream signaling events (Frooninckx *et al*. 2012). However, the specific neuropeptides and the GPCRs functioning in SPHK-1-mediated UPR^mt^ activation have not been identified.

FSHR-1 is a GPCR containing extracellular Leucine Rich Repeats (LRRs) homologous to the follicle-stimulating hormone receptor (Powell *et al*. 2009). FSHR-1 plays a critical role in activating innate immunity in response to infection by pathogenic bacteria, and functions in the intestine to promote protection to pathogenic infection and to regulate anti-microbial gene expression. (Cho *et al*. 2007; Powell *et al*. 2009; Miller *et al*. 2015). Interestingly, infection by pathogenic bacteria leads to multiple cellular responses in the intestine, including the activation of the UPR^mt^ (Liu *et al*. 2014; Pellegrino *et al*. 2014).

In this study, through the analysis of *fshr-1* null mutants, we show that FSHR-1 positively regulates UPR^mt^ activation in intestine and promotes the mitochondrial association of SPHK-1. Through tissue-specific rescue experiments, we find that FSHR-1 functions primarily in the nervous system, and also in the intestine, to exert its function in UPR^mt^ activation. We propose that FSHR-1 is part of a neuroendocrine signaling network that functions to activate the UPR^mt^ through inter-tissue signaling.

## Materials and Methods

### *C. elegans* strains

Strains used in this study were maintained at 22°C following standard methods. Young adult hermaphrodites derived from the wild type reference strain N2 Bristol were used for all experiments. The following mutant strains were used. SJ4100: *zcIs13[Phsp-6::GFP]*, OJ4113*: vjIs138[Pges-1::sphk-1::gfp], OJ4143: vjIs148[Pges-1::tomm-20::mCherry]*, OJ2329*: vjIs208[Pgst-4::gfp]*, OJ997 *sphk-1(ok1097)*, KP3397: *fshr-1(ok778)*. The *sphk-1(ok1097)* and *fshr-1(ok778)* strains were outcrossed at least 6 times with wild type animals prior to analysis.

### Molecular biology

*fshr-1* cDNA was cloned from *C. elegans* wild type cDNA and then inserted into the pPD49.26 expression vector using standard molecular biology techniques. The following plasmids were generated *pSK55[Pges-1::fshr-1], pSK56[Prab-3::fshr-1]*. Sequence files of plasmids are available upon request.

### Transgenic lines

Transgenic strains were generated by injecting expression constructs (10–25 ng/ul) and the co-injection marker; *pJQ70 [Pofm-1::mCherry, 40 ng/ul]*, *KP#708[Pttx-3::rfp, 40 ng/ul]* or *KP#1106 [Pmyo-2::gfp, 10 ng/ul]* into N2 or indicated mutants using standard techniques (Mello *et al*. 1991). At least three lines for each transgene were tested and a representative transgene was used for further experiments. The following transgenic lines were made: *vjEx1448[Prab-3::fshr-1], vjEx1449[Pges-1::fshr-1]*

### Microscopy and analysis

Fluorescence microscopy experiments were performed following previous methods (Kim and Sieburth 2018a). Briefly, L4 stage or young adult worms were immobilized by using 2,3-butanedione monoxime (BDM, 30 mg/mL; Sigma) in M9 buffer then mounted on 2% agarose pads for imaging. To quantify the fluorescence intensity of P*hsp-6::GFP* or P*gst-4::GFP*, *Z* stacks of the intestine posterior to the vulva were selected as a representative region because of low basal expression in the absence of stress. Images were captured with the Nikon eclipse 90i microscope equipped with a Nikon PlanApo 40 x or 60x or 100x objective (NA = 1.4) and a PhotometricsCoolsnap ES2 or a Hamamatsu Orca Flash LT+ CMOS camera.

Metamorph 7.0 software (Universal Imaging/Molecular Devices) was used to capture serial image stacks, and the maximum intensity was measured (Kim and Sieburth 2018b). Intensity quantification analysis was performed on the same day to equalize the absolute fluorescence levels between samples within same experimental set.

### RNA Interference

Feeding RNAi knockdown assay was performed following established protocols (Kamath and Ahringer 2003). Briefly, gravid adult animals were placed on RNAi plates seeded with HT115(DE3) bacteria transformed with L4440 vector containing a genomic fragment of the gene to be knocked down (or empty L4440 vector as a control), to collect eggs then removed after 4 hours to obtain an age-matched synchronized worm population. Young adult animals were used for subsequent assays.

### Stress induction assays

For drug-induced stress, transgenic L4 animals were transferred to fresh NGM plates seeded with HB101 bacteria, then 80 ul of stock solutions dissolved in M9 buffer of paraquat were added to plates for a final concentration of 0.4 mM paraquat. 24 hours later, adults were selected for fluorescence microscopy analysis. For imaging of animals that had been subjected to RNAi-induced knockdown, L4 animals grown on RNAi plates were transferred to new RNAi plates to obtain synchronized animals and imaged 24 hours later. For P*gst-4::GFP* imaging, young adult animals are incubated with 5mM arsenite or 50mM paraquat in liquid solution (in M9 buffer) for 1 hour then drug was washed out and worms were transferred onto new NGM plates for 4 hours before imaging. M9 buffer was used as a control for paraquat and arsenite treatment.

For paraquat survival assays, young adult animals were placed onto NGM plates containing 10mM paraquat for 17 hours. After 17 hours, the percentage of surviving animals was counted every 3 hours over the course of 15 hours. Survival assay were done in experimental duplicate for each biological duplicate.

### Statistical Analysis

Student’s t test (two-tailed) was used to determine the statistical significance. P values less than 0.01 or 0.001 are indicated with asterisks **(p< 0.01), ***(p<0.001), respectively. Error bars in the figures indicate the standard error of the mean (±S.E.M). The exact numbers of sample size (n) are indicated in each figure.

### Data Availability

Strains and plasmids used in this study are available upon request. The authors confirm that all data necessary for confirming the conclusions of the findings are present within the article, figures.

## Results

### FSHR-1/GPCR signaling regulates the UPR^mt^

Prior studies found that *fshr-1* mutants have normal lifespans (Powell *et al*. 2009), but exhibit enhanced sensitivity to lethality caused by the mitochondrial toxin and UPR^mt^ activator, paraquat (Miller *et al*. 2015). Because impairing UPR^mt^ activation causes sensitivity to paraquat-induced lethality (Nargund *et al*. 2012; Gatsi *et al*. 2014; Liu *et al*. 2014; Kim and Sieburth 2018a), we speculated that FSHR-1 may positively regulate the UPR^mt^. To monitor UPR^mt^ activation, we quantified intestinal fluorescence of the UPR^mt^ transcriptional reporter, *zcIs13*, in which GFP is expressed under control of the *hsp-6* promoter (P*hsp-6::GFP*). Paraquat is an oxidant that interferes with electron transport at the inner mitochondrial membrane, and has been used widely to acutely activate the UPR^mt^ in both *C. elegans* and in mammalian cells (Nargund *et al*. 2012; Runkel *et al*. 2013; Fiorese *et al*. 2016; Kim and Sieburth 2018a). Wild type animals treated with paraquat for 24 hours exhibited a greater than ten-fold increase in P*hsp-6::GFP* expression in the intestine compared to non-treated controls. In contrast, *fshr-1* mutants showed either no significant change or a small increase in P*hsp-6::GFP* expression following paraquat treatment (Figure 1A and 2A). *cco-1* (aka *cox-5B*), encodes the cytochrome *c* oxidase subunit in complex IV, which is the terminal electron acceptor of the electron transport chain, and RNA interference (RNAi)-mediated *cco-1* knockdown is a potent activator of the UPR^mt^ (Nargund *et al*. 2012; Berendzen *et al*. 2016; Merkwirth *et al*. 2016; TIAN *et al*. 2016; Kim and Sieburth 2018a). *cco-1* RNAi by feeding significantly increased P*hsp-6::GFP* expression in wild type animals but failed to increase P*hsp-6::GFP* expression in *fshr-1* mutants (Figure 1C). *fshr-1* mutants exhibited normal induction of the antioxidant reporter P*gst-4::GFP* (Inoue *et al*. 2005; Choe *et al*. 2009; Ruiz-Ramos *et al*. 2009; Prakash *et al*. 2015; Wu *et al*. 2016) in response to treatment with either the mitochondrial ROS generator arsenite, or with paraquat (Figure 1D). Thus, FSHR-1 positively regulates UPR^mt^ activation in response to mitochondrial stress.

**Figure 1.**
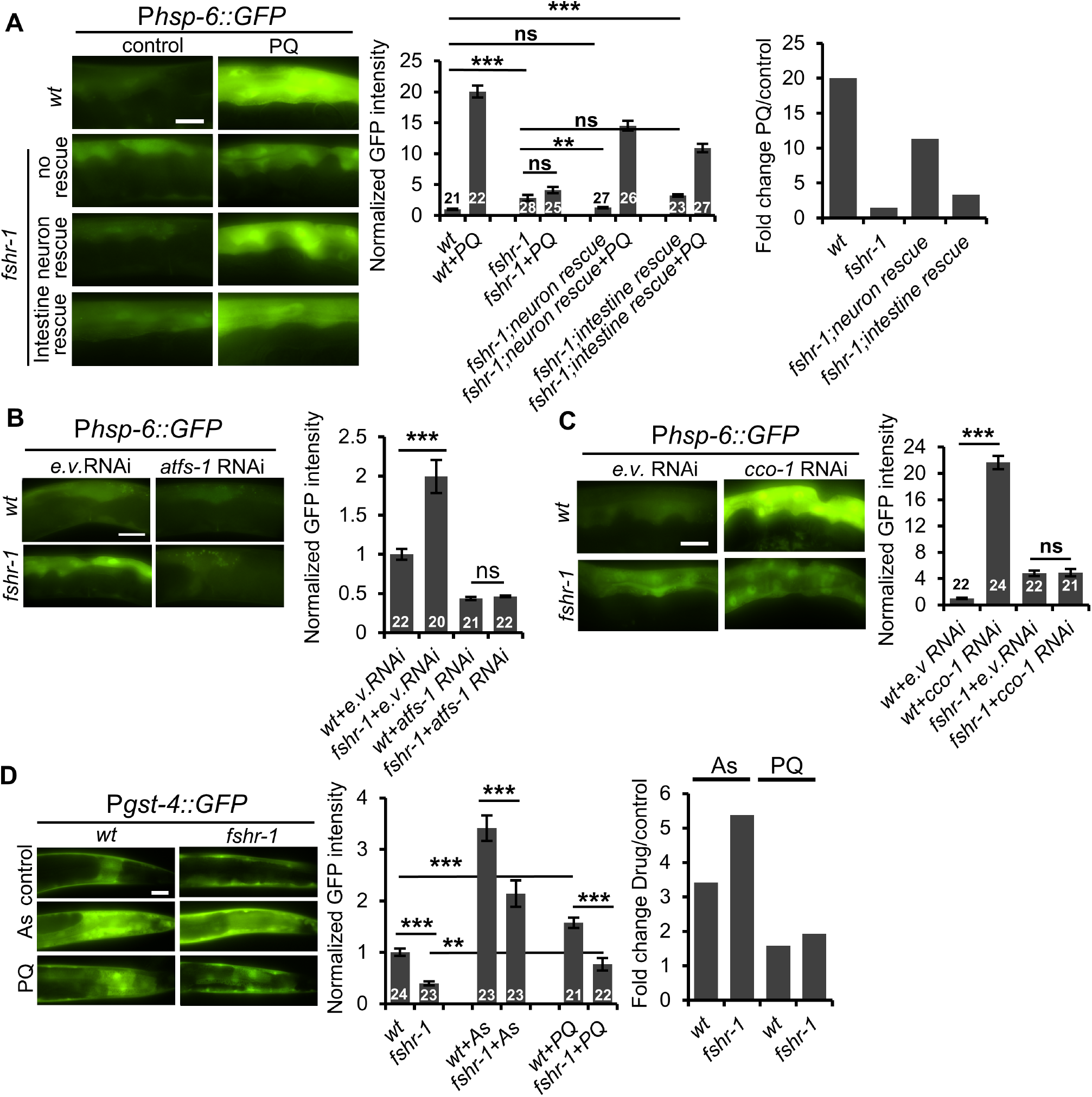
FSHR-1 functions in the nervous system to activate the UPR^mt^. (A) *Left*, Representative images of posterior intestines of young adult animals or *fshr-1* mutants expressing the *zcIs13* [P*hsp-6::GFP*] transgene (*GFP* driven by the *hsp-6* promoter) following treatment with M9 (control) or paraquat (PQ) for 24 hours. “*fshr-1* neuronal rescue” denotes *fshr-1* mutants expressing *fshr-1* cDNA under the neuron-specific promoter, *rab-3*. “*fshr-1* intestine rescue” denotes *fshr-1* mutants expressing *fshr-1* cDNA under the intestine-specific promoter, *ges-1. Middle*, Average GFP fluorescence intensity is quantified. *Right*, Fold changes of P*hsp-6::GFP* expression of the indicated strains by paraquat treatment are measured. (B) *Left*, Representative images of intestines of wild type or *fshr-1* mutants expressing P*hsp-6::GFP* following empty vector (e.v.) control or *atfs-1* RNAi treatment. *Right*, Average GFP fluorescence intensity is quantified. (C) *Left*, Representative images of P*hsp-6::GFP* fluorescence in intestines of wild type or *fshr-1* mutants following empty vector (e.v.) control or *cco-1/cox-5B* RNAi treatment. *Right*, Average GFP fluorescence intensity is quantified. (D) *Left*, Representative images of posterior intestines from animals expressing the antioxidant reporter transgene [P*gst-4::GFP*] in wild type or *fshr-1* mutants in the absence or presence of arsenite (As), or paraquat (PQ). *Middle*, Average GFP fluorescence intensity is quantified. *Right*, Fold changes of *gst-4* expression of indicated strains by arsenite or paraquat treatment are measured. Scale bars represent 10 μm. Error bars indicate ±S.E.M. The number of animals tested are indicated. Student’s t-test **p<0.01, ***p<0.001. “ns” denotes “not statistically significant”

**Figure 2.**
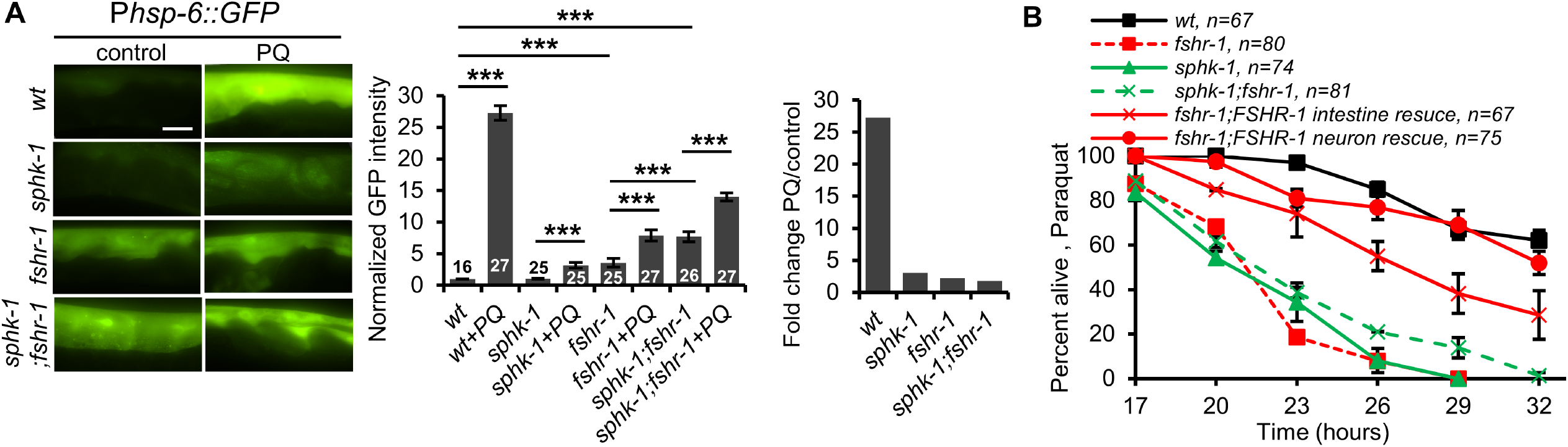
FSHR-1 functions in a common pathway with SPHK-1 to activate the UPR^mt^. (A) *Left*, Representative images of P*hsp-6::GFP* expression in the intestines of wild type, *sphk-1* mutants, *fshr-1* mutants, and *sphk-1;fshr-1* double mutants in the absence or the presence of paraquat (PQ) for 24 hours. *Middle*, Average GFP fluorescence intensity is quantified. *Right*, Fold change denotes the ratio of average fluorescence intensity of animals treated with paraquat (PQ) and control animals treated with buffer (M9). (B) Survival rate curves of the indicated strains on plates containing 10mM paraquat. The number of animals tested are indicated. Scale bar represents 10 μm. Student’s t-test ***p<0.001.

### FSHR-1/GPCR functions cell non-autonomously in neurons to regulate UPR^mt^

FSHR-1 is expressed primarily in the intestine and in a subset of neurons (Sieburth *et al*. 2005; Cho *et al*. 2007). To determine in which tissue FSHR-1 functions to regulate the UPR^mt^, we examined P*hsp-6::GFP* expression in transgenic *fshr-1* mutants expressing *fshr-1* cDNA in either the nervous system (using the *rab-3* promoter) or the intestine (using the *ges-1* promoter). Pan-neuronal *fshr-1* nearly fully rescued the paraquat-induced *hsp-6* expression defects of *fshr-1* mutants. In contrast, intestinal *fshr-1* expression only partially rescued the paraquat-induced *hsp-6* expression defects of *fshr-1* mutants (Figure 1A). P*hsp-6* expression was increased 11 fold by pan-neuronal *fshr-1* compared to just 3.3 fold by intestinal *fshr-1* expression (Figure 1A). Taken together, these results suggest that FSHR-1 primarily functions cell non-autonomously in the nervous system to positively regulate UPR^mt^ activation, and intestinal *fshr-1* has a minor role in contributing to UPR^mt^ activation.

### Neuronal FSHR-1 regulates baseline *hsp-6* expression

Under non-stressed conditions, *fshr-1* mutants exhibited a small but significant increase in P*hsp-6::GFP* expression in the intestine compared to wild type controls (Figure 1A and 1C). The increase in P*hsp-6::GFP* expression was blocked by RNAi-mediated knockdown of *atfs-1* (Figure 1B), suggesting that FSHR-1 keeps UPR^mt^ activation low under non-stressed conditions. The increase in baseline P*hsp-6::GFP* expression of *fshr-1* mutants was restored to wild type levels by pan-neuronal, but not by intestinal *fshr-1* cDNA expression (Figure 1A). Thus, neuronal FSHR-1 functions cell non-autonomously to keep UPR^mt^ activity low in the absence of stress.

### FSHR-1 and SPHK-1 function in a common pathway to activate the UPR^mt^

SPHK-1 functions in the intestine to activate the UPR^mt^ in response to a variety of mitochondrial stressors, including paraquat (Kim and Sieburth 2018a). To test whether FSHR-1 functions in a common pathway with SPHK-1 to activate the UPR^mt^, we examined genetic interactions between *sphk-1* and *fshr-1* mutants. *sphk-1* mutants exhibit a significant reduction in paraquat-induced expression of P*hsp-6::GFP*, (89% reduction) that is similar to that exhibited by *fshr-1* mutants (92% reduction (Figure 2A and (Kim and Sieburth 2018a)). Double mutants lacking both *sphk-1* and *fshr-1* displayed defects in paraquat-induced *hsp-6* induction that were no more severe than those seen in single mutants (93% reduction, Figure 2A).

*sphk-1* mutants exhibit reduced survival rates when exposed to toxic levels of paraquat (Kim and Sieburth 2018a), that are similar to those of *fshr-1* mutants (Figure 2B). The survival rates of double mutants lacking both *sphk-1* and *fshr-1* were no more severe than those of either single mutant (Figure 2B). Expression of *sphk-1* cDNA in the intestine fully rescued the increased paraquat-induced lethality of *sphk-1* mutants (Kim and Sieburth 2018a). In contrast, expression of *fshr-1* cDNA in the nervous system fully rescued the increased paraquat-induced lethality of *fshr-1* mutants. In addition, expression of *fshr-1* cDNA in the intestine restored near-normal paraquat sensitivity to *fshr-1* mutants (Figure 2B and (Miller *et al*. 2015)). Taken together, FSHR-1 and SPHK-1 function in a common pathway to activate the UPR^mt^ and to promote organism-wide protection from mitochondrial stress-induced lethality. Moreover, FSHR-1 functions in both the nervous system and the intestine to activate the UPR^mt^ and promote survival, whereas SPHK-1 functions exclusively in the intestine.

### FSHR-1/GPCR regulates mitochondrial association of SPHK-1 in the intestine

SPHK-1 rapidly associates with intestinal mitochondria following mitochondrial stress (Kim and Sieburth 2018a). To determine whether FSHR-1 regulates stress-induced SPHK-1 mitochondrial association, we examined the mitochondrial abundance of functional SPHK-1::GFP fusion proteins before and after paraquat exposure. Wild type animals treated with paraquat exhibit a 1.8 fold increase in SPHK-1::GFP mitochondrial fluorescence intensity (Figure 3C and (Kim and Sieburth 2018a)). *fshr-1* mutants exhibited three defects in SPHK-1::GFP mitochondrial association. First, *fshr-1* mutants exhibited a slightly smaller 1.5 fold increase in paraquat-induced mitochondrial SPHK-1::GFP fluorescence compared to wild type controls (Figure 3C). Second, *fshr-1* mutants exhibited a nearly two-fold reduction in SPHK-1::GFP mitochondrial fluorescence compared to wild type controls in the absence of paraquat (Figure 3A). Third, *fshr-1* mutants exhibited a nearly two-fold reduction in SPHK-1::GFP mitochondrial fluorescence compared to wild type controls following paraquat treatment (Figure 3C). The reduction in SPHK-1::GFP mitochondrial fluorescence is likely not due to decreased expression of the *sphk-1::gfp* transgene, or decreased mitochondrial mass since the fluorescence intensity of the mitochondrial marker TOMM-20::mCherry expressed under the same promoter (*ges-1*) was not reduced in *fshr-1* mutants either in the absence or presence of paraquat (Figure 3A and (Kim and Sieburth 2018a)). Thus, *fshr-1* mutations impair SPHK-1 association with mitochondria under both stressed and non-stressed conditions.

**Figure 3.**
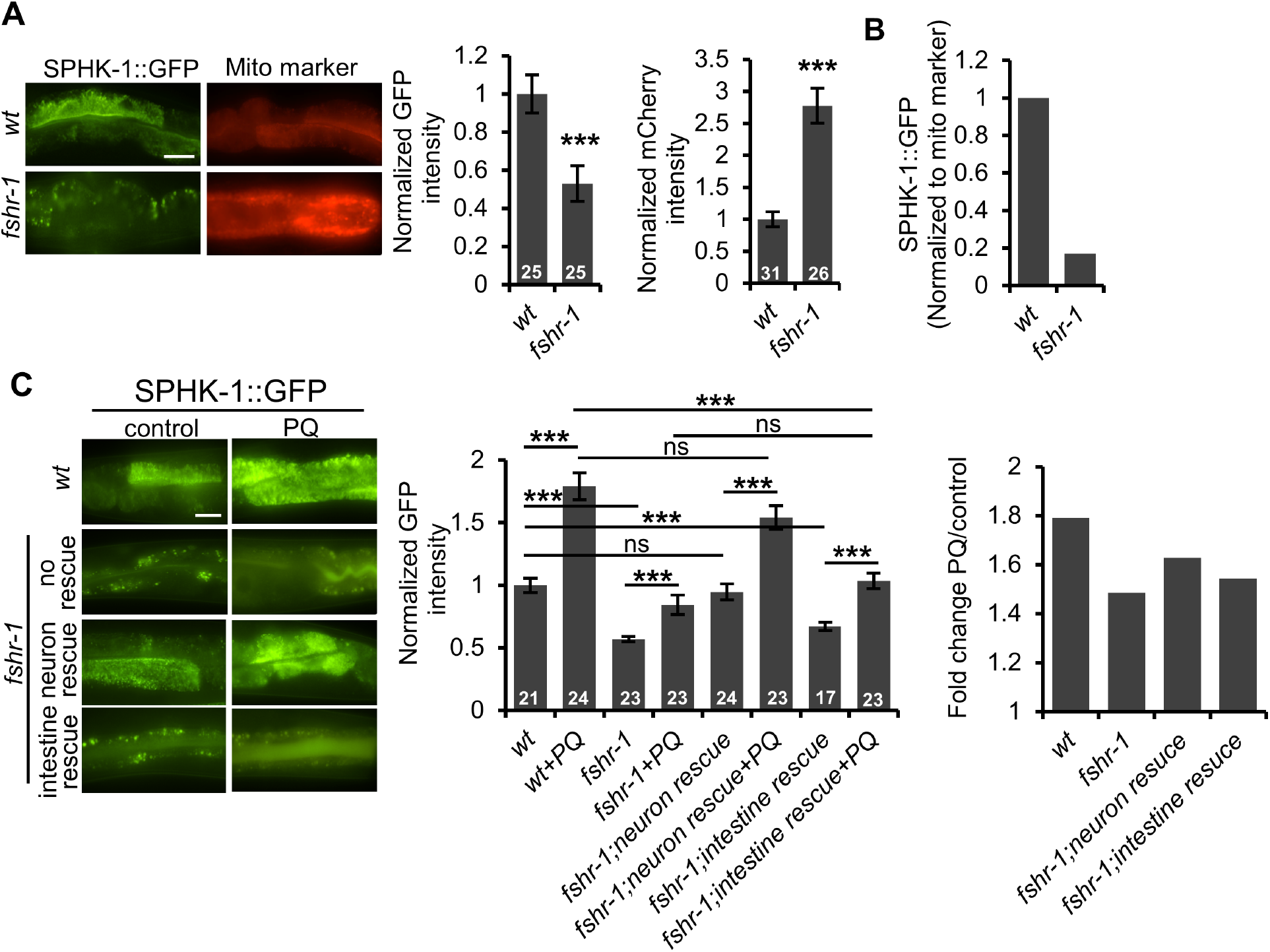
FSHR-1 regulates mitochondrial association of SPHK-1. (A) *Left*, Representative posterior intestine images of wild type or *fshr-1* mutants expressing SPHK-1::GFP or the mito-marker TOMM-20::mCherry under the intestinal-specific promoter, *ges-1*. *Right*, Average GFP and mCherry fluorescence intensities are quantified. (B) Ratio of SPHK-1::GFP expression in wild type or *fshr-1* mutants (normalized to the intensity of each of the mitochondria markers). (C) *Left*, Representative images of the indicated mutants expressing intestinal SPHK-1::GFP in the absence or presence of paraquat for 24 hours. *fshr-1* neuron, intestine rescue denotes transgenic *fshr-1* mutants expressing *fshr-1* cDNA under the pan-neural promoter, *rab-3*, or the intestinal promoter, *ges-1*, respectively. *Middle*, Average fluorescence intensities in the indicated strains expressing SPHK-1::GFP are quantified. *Right*, Fold changes of SPHK-1::GFP expression of the indicated strains by paraquat treatment are measured. Scale bar represents 10 μm. The number of animals tested are indicated. Student’s t-test ***p<0.001. “ns” denotes “not statistically significant”

Interestingly, we observed an increase in mitochondrial mass in *fshr-1* mutants, as revealed by an increase in TOMM-20::mCherry mitochondrial fluorescence compared to wild type controls (Figure 3A). Factoring in the increased mitochondrial mass in *fshr-1* mutants, we found an approximately 5.9 fold decrease in mitochondrial-associated SPHK-1 at baseline (Figure 3B). Thus, FSHR-1 negatively regulates mitochondrial mass in the intestine and positively regulates SPHK-1 mitochondrial association, under normal conditions and following stress.

### Neuronal *fshr-1* cell non-autonomously regulates mitochondrial association of SPHK-1 in intestine

To determine in which tissue FSHR-1 functions to regulate the mitochondrial association of SPHK-1, we quantified SPHK-1::GFP mitochondrial fluorescence in *fshr-1* mutants expressing *fshr-1* cDNA in either the nervous system or in the intestine. We found that pan-neuronal expression of *fshr-1* cDNA in *fshr-1* mutants restored mitochondrial SPHK-1::GFP fluorescence intensity to wild type levels, both in the absence of paraquat and following paraquat treatment. In contrast, intestinal *fshr-1* cDNA expression failed to rescue the SPHK-1::GFP fluorescence defects of *fshr-1* mutants in the absence of paraquat and partially restored paraquat-induced SPHK-1 mitochondrial association (Figure 3C). These results suggest that FSHR-1 functions exclusively in the nervous system to regulate SPHK-1 abundance in the intestine in the absence of stress and that FSHR-1 function in both in the nervous system and in the intestine is likely to promote paraquat-induced SPHK-1 mitochondrial association.

## Discussion

The nervous system coordinates various stress responses by releasing diffusible factors that act upon distal tissues to activate cellular defense programs (Berendzen *et al*. 2016; Shao *et al*. 2016). Here we show that the conserved GPCR FSHR-1 is part of an integrated organism-wide response to mitochondrial stress that functions to activate the UPR^mt^ in the intestine. FSHR-1 promotes UPR^mt^ activation in response to either acute exposure to mitochondrial toxins or to chronic mitochondrial dysfunction. FSHR-1 functions in a common pathway with SPHK-1 to promote survival in response to toxic mitochondrial stress, and to promote UPR^mt^ activation. FSHR-1 positively regulates the mitochondrial association of SPHK-1 in the intestine in the absence of stress, as well as stress-induced SPHK-1 mitochondrial recruitment. FSHR-1 functions in the nervous system and in the intestine to promote UPR^mt^ activation, stress-induced SPHK-1 mitochondrial recruitment and survival in the presence of mitochondrial stress. We propose a model whereby FSHR-1 activates the UPR^mt^ by two mechanisms: 1) neuronal FSHR-1 promotes cell non-autonomous activation of the UPR^mt^ by regulating intestinal SPHK-1 activity, and 2) intestinal FSHR-1 activates the UPR^mt^ by a cell-autonomous mechanism that does not involve SPHK-1 (Figure 4). FSHR-1 also regulates mitochondrial homeostasis in the absence of stress by regulating mitochondrial mass, baseline UPR^mt^ activity, and SPHK-1 mitochondrial association.

**Figure 4.**
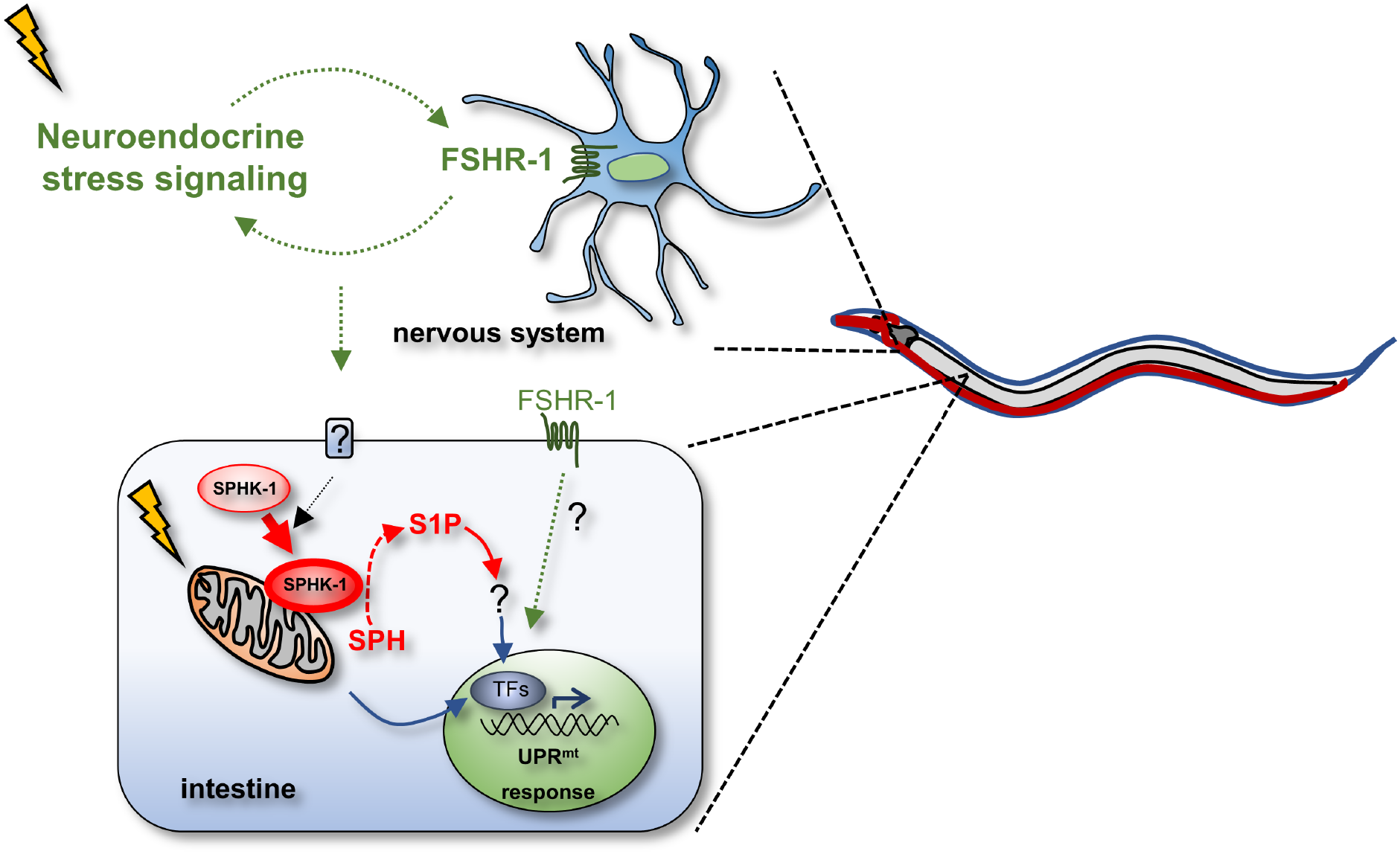
Working model for UPR^mt^ activation by FSHR-1 and SPHK-1. Upon mitochondrial stress, neuroendocrine signaling involving FSHR-1 signaling from the nervous system activates the UPR^mt^ by regulating intestinal SPHK-1 abundance on mitochondria, where SPHK-1 catalyzes the production of S1P from SPH. S1P then activates the UPR^mt^ transcriptional response. FSHR-1 also functions cell autonomously in the intestine to promote activation of the UPR^mt^ by a mechanism that may be independent of SPHK-1.

Previous studies have established an important role for FSHR-1 in activating the innate immune response following infection with pathogenic bacteria. FSHR-1 promotes both survival following infection, and the behavioral avoidance response to pathogenic bacteria (Powell *et al*. 2009; Miller *et al*. 2015), and FSHR-1 activates a number of antimicrobial and antioxidant genes in response to pathogens (Powell *et al*. 2009; Miller *et al*. 2015). FSHR-1 is proposed to not participate in detection of the pathogenic bacteria themselves, but instead to participate in the detection of cellular damage (e.g. ROS) resulting from bacterial infection (Miller *et al*. 2015). Pathogenic bacterial infection is also a potent activator of the UPR^mt^, and UPR^mt^ activation by pathogens promotes survival by inducing the expression of innate immune genes (Pellegrino *et al*. 2014). Thus, FSHR-1 may activate the innate immune response by contributing to activation of the UPR^mt^ following infection. Interestingly, prior studies have found that expression of FSHR-1 in the nervous system or the intestine can rescue survival defects of *fshr-1* mutants (Powell *et al*. 2009), consistent with our studies showing a function for FSHR-1 in UPR^mt^ activation in either tissue. We speculate that FSHR-1 may be a central component in the cellular response to mitochondrial dysfunction caused by a broad array of insults (e.g. pathogenic infection, ROS, and UPR^mt^ dysfunction).

FSHR-1 also regulates intestinal mitochondrial homeostasis in the absence of stress. *fshr-1* mutants exhibit a nearly three-fold increase in mitochondrial mass in the intestine, and a corresponding reduction in mitochondrial-associated SPHK-1. FSHR signaling negatively regulates mitochondrial biogenesis in mammals (Liu *et al*. 2017). Enhanced mitochondria biogenesis is correlated with UPR^mt^ activation. Inducers of mitochondria biogenesis, such as nicotinamide riboside (NR) and poly(ADP-ribose) polymerase inhibitors (PARPi) disrupt the mito-nuclear protein homeostasis resulting in UPR^mt^ activation via the sirtuin 1 (SIRT1) pathway in mammals and *C. elegans*. (Mouchiroud *et al*. 2013). In addition, rapamycin and resveratrol, which increase mitochondrial content, also induce UPR^mt^ activation in *C. elegans* (Ungvari *et al*. 2011; Houtkooper *et al*. 2013; Lerner *et al*. 2013). Consistent with this, *fshr-1* mutants exhibit increased baseline *hsp-6* expression, which is dependent upon *atfs-1*, suggesting a potential function for FSHR-1 in suppressing UPR^mt^ activation by reducing mitochondrial mass and/or biogenesis. We speculate that the defects in paraquat-induced UPR^mt^ activation in *fshr-1* mutants may be due to an underlying defect in the mitochondrial association of SPHK-1 under normal conditions.

Since FSHR-1 functions to protect animals from diverse stressors, the ligand for FSHR-1 is likely to originate from the host rather than from an exogenous source (Powell *et al*. 2009). FSHR-1 shares homology with three human GPCRs: FSHR, TSHR and LHCGR by virtue of nine LRRs found in its extracellular domain (Dolan *et al*. 2007). These GPCRs are activated by the heterodimeric glycopeptide hormones FSHα/β, TSHα/β, and LHα/β, respectively. In humans, FSH induces the generation of S1P by stimulating SphK1 activity for granulosa cell proliferation (Hernandez-Coronado *et al*. 2016), suggesting a potential functional conservation of ligand-activated FSHR-1 signaling. However, the worm genome does not encode obvious orthologues of any of these ligands (Powell *et al*. 2009; Miller *et al*. 2015), suggesting divergence of an ancestral FSHR-1 ligand. A number of GPCR ligands have been identified that function non-autonomously to either activate the UPR^mt^ or the innate immunity response (Shao *et al*. 2016; Kim and Ewbank 2018). Among these are a set of neuropeptide-like proteins that function in a subset of sensory/interneurons to activate the UPR^mt^. These peptides are proposed to function in a neuronal circuit that transduces mitochondrial stress signals originating in the nervous system to the intestine to activate the UPR^mt^ (Shao *et al*. 2016). Because of the requirement for FSHR-1 in the nervous system in activating the UPR^mt^, we propose that FSHR-1 functions as part of this neuroendocrine signaling network, possibly as a receptor for one or more of these peptides. Consistent with a role for FSHR-1 in regulating neuron function, FSHR-1 positively regulates synaptic transmission at motor synapses (Sieburth *et al*. 2005). Identifying the ligand(s) for FSHR-1 will help to clarify both the cell autonomous and cell non-autonomous mechanisms by which FSHR-1 activates the UPR^mt^ pathway.

## Acknowledgements

This work was supported by grants from the NIH National institute of Neurological Disorders and Stroke (NINDS) to D.S. (NS071085 and NS099414). Some strains were provided by the Caenorhabditis Genetics Center (CGC), which is funded by the NIH office of Research infrastructure Programs (P40 OD010440).

## References

Berendzen, K. M., J. Durieux, L. W. Shao, Y. Tian, H. E. Kim et al., 2016 Neuroendocrine Coordination of Mitochondrial Stress Signaling and Proteostasis. Cell 166: 1553–1563 e1510.

Cho, S., K. W. Rogers and D. S. Fay, 2007 The C. elegans glycopeptide hormone receptor ortholog, FSHR-1, regulates germline differentiation and survival. Curr Biol 17: 203–212.

Choe, K. P., A. J. Przybysz and K. Strange, 2009 The WD40 repeat protein WDR-23 functions with the CUL4/DDB1 ubiquitin ligase to regulate nuclear abundance and activity of SKN-1 in Caenorhabditis elegans. Mol Cell Biol 29: 2704–2715.

Dolan, J., K. Walshe, S. Alsbury, K. Hokamp, S. O’Keeffe et al., 2007 The extracellular leucine-rich repeat superfamily; a comparative survey and analysis of evolutionary relationships and expression patterns. BMC Genomics 8: 320.

Durieux, J., S. Wolff and A. Dillin, 2011 The cell-non-autonomous nature of electron transport chain-mediated longevity. Cell 144: 79–91.

Fiorese, C. J., A. M. Schulz, Y. F. Lin, N. Rosin, M. W. Pellegrino et al., 2016 The Transcription Factor ATF5 Mediates a Mammalian Mitochondrial UPR. Curr Biol 26: 2037–2043.

Frooninckx, L., L. Van Rompay, L. Temmerman, E. Van Sinay, I. Beets et al., 2012 Neuropeptide GPCRs in C. elegans. Front Endocrinol (Lausanne) 3: 167.

Gatsi, R., B. Schulze, M. J. Rodriguez-Palero, B. Hernando-Rodriguez, R. Baumeister et al., 2014 Prohibitin-mediated lifespan and mitochondrial stress implicate SGK-1, insulin/IGF and mTORC2 in C. elegans. PLoS One 9: e107671.

Hernandez-Coronado, C. G., A. Guzman, A. Rodriguez, J. A. Mondragon, M. C. Romano et al., 2016 Sphingosine-1-phosphate, regulated by FSH and VEGF, stimulates granulosa cell proliferation. Gen Comp Endocrinol 236: 1–8.

Houtkooper, R. H., L. Mouchiroud, D. Ryu, N. Moullan, E. Katsyuba et al., 2013 Mitonuclear protein imbalance as a conserved longevity mechanism. Nature 497: 451–457.

Inoue, H., N. Hisamoto, J. H. An, R. P. Oliveira, E. Nishida et al., 2005 The C. elegans p38 MAPK pathway regulates nuclear localization of the transcription factor SKN-1 in oxidative stress response. Genes Dev 19: 2278–2283.

Kamath, R. S., and J. Ahringer, 2003 Genome-wide RNAi screening in Caenorhabditis elegans. Methods 30: 313–321.

Kim, D. H., and J. J. Ewbank, 2018 Signaling in the innate immune response. WormBook 2018: 1–35.

Kim, S., and D. Sieburth, 2018a Sphingosine Kinase Activates the Mitochondrial Unfolded Protein Response and Is Targeted to Mitochondria by Stress. Cell Rep 24: 2932–2945 e2934.

Kim, S., and D. Sieburth, 2018b Sphingosine Kinase Regulates Neuropeptide Secretion During the Oxidative Stress-Response Through Intertissue Signaling. J Neurosci 38: 8160–8176.

Lerner, C., A. Bitto, D. Pulliam, T. Nacarelli, M. Konigsberg et al., 2013 Reduced mammalian target of rapamycin activity facilitates mitochondrial retrograde signaling and increases life span in normal human fibroblasts. Aging Cell 12: 966–977.

Liu, P., Y. Ji, T. Yuen, E. Rendina-Ruedy, V. E. DeMambro et al., 2017 Blocking FSH induces thermogenic adipose tissue and reduces body fat. Nature 546: 107–112.

Liu, Y., B. S. Samuel, P. C. Breen and G. Ruvkun, 2014 Caenorhabditis elegans pathways that surveil and defend mitochondria. Nature 508: 406–410.

Martinez, B. A., D. A. Petersen, A. L. Gaeta, S. P. Stanley, G. A. Caldwell et al., 2017 Dysregulation of the Mitochondrial Unfolded Protein Response Induces Non-Apoptotic Dopaminergic Neurodegeneration in C. elegans Models of Parkinson’s Disease. J Neurosci 37: 11085–11100.

Mello, C. C., J. M. Kramer, D. Stinchcomb and V. Ambros, 1991 Efficient gene transfer in C.elegans: extrachromosomal maintenance and integration of transforming sequences. EMBO J 10: 3959–3970.

Merkwirth, C., V. Jovaisaite, J. Durieux, O. Matilainen, S. D. Jordan et al., 2016 Two Conserved Histone Demethylases Regulate Mitochondrial Stress-Induced Longevity. Cell 165: 1209–1223.

Miller, E. V., L. N. Grandi, J. A. Giannini, J. D. Robinson and J. R. Powell, 2015 The Conserved G-Protein Coupled Receptor FSHR-1 Regulates Protective Host Responses to Infection and Oxidative Stress. PLoS One 10: e0137403.

Mouchiroud, L., R. H. Houtkooper, N. Moullan, E. Katsyuba, D. Ryu et al., 2013 The NAD(+)/Sirtuin Pathway Modulates Longevity through Activation of Mitochondrial UPR and FOXO Signaling. Cell 154: 430–441.

Nargund, A. M., C. J. Fiorese, M. W. Pellegrino, P. Deng and C. M. Haynes, 2015 Mitochondrial and nuclear accumulation of the transcription factor ATFS-1 promotes OXPHOS recovery during the UPR(mt). Mol Cell 58: 123–133.

Nargund, A. M., M. W. Pellegrino, C. J. Fiorese, B. M. Baker and C. M. Haynes, 2012 Mitochondrial import efficiency of ATFS-1 regulates mitochondrial UPR activation. Science 337: 587–590.

Pellegrino, M. W., A. M. Nargund, N. V. Kirienko, R. Gillis, C. J. Fiorese et al., 2014 Mitochondrial UPR-regulated innate immunity provides resistance to pathogen infection. Nature 516: 414–417.

Powell, J. R., D. H. Kim and F. M. Ausubel, 2009 The G protein-coupled receptor FSHR-1 is required for the Caenorhabditis elegans innate immune response. Proc Natl Acad Sci U S A 106: 2782–2787.

Prakash, C., M. Soni and V. Kumar, 2015 Biochemical and Molecular Alterations Following Arsenic-Induced Oxidative Stress and Mitochondrial Dysfunction in Rat Brain. Biol Trace Elem Res 167: 121–129.

Ruiz-Ramos, R., L. Lopez-Carrillo, A. D. Rios-Perez, A. De Vizcaya-Ruiz and M. E. Cebrian, 2009 Sodium arsenite induces ROS generation, DNA oxidative damage, HO-1 and c-Myc proteins, NF-kappaB activation and cell proliferation in human breast cancer MCF-7 cells. Mutat Res 674: 109–115.

Runkel, E. D., S. Liu, R. Baumeister and E. Schulze, 2013 Surveillance-activated defenses block the ROS-induced mitochondrial unfolded protein response. PLoS Genet 9: e1003346.

Shao, L. W., R. Niu and Y. Liu, 2016 Neuropeptide signals cell non-autonomous mitochondrial unfolded protein response. Cell Res 26: 1182–1196.

Sieburth, D., Q. Ch’ng, M. Dybbs, M. Tavazoie, S. Kennedy et al., 2005 Systematic analysis of genes required for synapse structure and function. Nature 436: 510–517.

Tian, Y., G. Garcia, Q. Bian, K. K. Steffen, L. Joe et al., 2016 Mitochondrial Stress Induces Chromatin Reorganization to Promote Longevity and UPR(mt). Cell 165: 1197–1208.

Ungvari, Z., W. E. Sonntag, R. de Cabo, J. A. Baur and A. Csiszar, 2011 Mitochondrial protection by resveratrol. Exerc Sport Sci Rev 39: 128–132.

Wu, C. W., A. Deonarine, A. Przybysz, K. Strange and K. P. Choe, 2016 The Skp1 Homologs SKR-1/2 Are Required for the Caenorhabditis elegans SKN-1 Antioxidant/Detoxification Response Independently of p38 MAPK. PLoS Genet 12: e1006361.

Zhang, Q., X. Wu, P. Chen, L. Liu, N. Xin et al., 2018 The Mitochondrial Unfolded Protein Response Is Mediated Cell-Non-autonomously by Retromer-Dependent Wnt Signaling. Cell 174: 870–883 e817.

